# Nigral pathology contributes to microstructural integrity of striatal and frontal tracts in Parkinson’s disease

**DOI:** 10.1101/2022.12.21.521411

**Authors:** Chen-Pei Lin, Lydian EJ Knoop, Irene Frigerio, John GJM Bol, Annemieke JM Rozemuller, Henk W Berendse, Petra JW Pouwels, Wilma DJ van de Berg, Laura E Jonkman

## Abstract

**Background:** Motor and cognitive impairment in Parkinson’s disease (PD) is associated with dopaminergic dysfunction that stems from substantia nigra (SN) degeneration and concomitant α-synuclein accumulation. Diffusion MRI can detect microstructural alterations of the SN and its tracts to (sub)cortical regions, but their pathological sensitivity is still poorly understood.

**Objective:** To unravel the pathological substrate underlying microstructural alterations of the SN, and its tracts to the dorsal striatum and dorsolateral prefrontal cortex (DLPFC) in PD.

**Methods:** Combining post-mortem *in-situ* MRI and histopathology, T1-weighted and diffusion MRI of 9 PD, 6 PD with dementia (PDD), 5 dementia with Lewy bodies (DLB), and 10 control donors were collected. From MRI, mean diffusivity (MD) and fractional anisotropy (FA) were derived from the SN, and tracts between the SN and caudate nucleus, putamen, and DLPFC. Phosphorylated-Ser129-α-synuclein and tyrosine hydroxylase immunohistochemistry was included to quantify nigral Lewy pathology and dopaminergic degeneration, respectively.

**Results:** Compared to controls, PD and PDD/DLB showed increased MD of the SN and SN-DLPFC tract, as well as increased FA of the SN-caudate nucleus tract. Both PD and PDD/DLB showed nigral Lewy pathology and dopaminergic loss compared to controls. Increased FA of the SN and SN-caudate nucleus tract was associated with SN dopaminergic loss, while increased MD of the SN-DLPFC tract was associated with increased SN Lewy neurite load.

**Conclusions:** In PD and PDD/DLB, diffusion MRI captures microstructural alterations of the SN and tracts to the dorsal striatum and DLPFC, which differentially associates with SN dopaminergic degeneration and Lewy neurite pathology.

## Introduction

Parkinson’s disease (PD) is the second most prevalent neurodegenerative disease to date, and manifests with a wide range of motor and cognitive symptoms^1–4^. Motor features, such as resting tremor, bradykinesia and rigidity, have been associated with reduced dopaminergic innervation to the striatum, as a consequence of dopaminergic loss within the substantia nigra (SN) and concomitant pathological a-syn accumulation in the form of Lewy bodies (LB) and Lewy neurites (LN). Cognitive function, particularly in attention, memory and executive domains, are commonly impaired in PD, and more pronouncedly dysfunctional in PD dementia (PDD). Up to 80% of PD patients develop PDD during the course of the disease^5–7^. In addition, patients that develop dementia before or within one year of motor symptoms onset are diagnosed as dementia with Lewy bodies (DLB). To predict and monitor disease progression in PD, accurate, reliable, and neuropathologically verified biomarkers are urgently needed^8–10^.

Diffusion MRI has great potential to assess microstructural alterations in neurological disorders^11–14^. In both early- and advanced-stage PD studies reveal increased fractional anisotropy (FA) and mean diffusivity (MD) of the SN and its tracts to the dorsal striatum. In turn, these microstructural alterations were correlated with motor severity on the unified Parkinson’s disease rating scale (UPDRS)^15–18^. Furthermore, a longitudinal study showed progressively increased MD in PD patients at follow-ups^19^, suggesting that diffusion MRI measures could be a potential indicator for monitoring disease progression^16,20^. Despite a large body of diffusion MRI studies in PD, the underlying pathological substrate of these diffusion changes is still unclear. Multi-modal imaging, combining dopaminergic transporter imaging (DAT) and MRI, attempted to unravel this but reported no correlation between dopaminergic loss in the putamen, and MD or FA of the nigrostriatal tract ^21^.

Cognitive dysfunction is highly prevalent in PD^22^, and serves as a predictor for progression to PDD^23–25^. The cortico-basal ganglia circuitry, which connects the cortex, basal ganglia and thalamus, provides a neuroanatomical substrate of cognitive processing. Particularly, the dorsolateral prefrontal cortex (DLPFC) is highly engaged in mediating attention and memory processing. In PD, widespread α-syn accumulation and dopaminergic depletion in the DLPFC, has shown to impair these functions^26–29^. However, the impact of nigral pathology on this tract remains elusive.

The current study aims to unravel the pathological substrate(s) of microstructural alterations in tracts between the SN, dorsal striatum and DLPFC in PD and PDD/DLB and age-matched controls, using a within-subject post-mortem *in situ* MRI and histopathology approach. We hypothesize that both PD and PDD/DLB show altered microstructure of the tracts between the SN and dorsal striatum, and that PDD/DLB show particularly an alteration in tract between the SN and DLPFC, due to its implication in cognitive dysfunction. We further hypothesize that these tract alterations correlate with pathology within the SN. This study may bridge the gap between neuroimaging and PD pathological features, and aid the development of pathology-sensitive MRI biomarkers that have the potential to predict and monitor motor and cognitive decline during disease progression.

## Methods

### Donor inclusion

A total of 30 clinically defined and pathologically confirmed brain donors were included in the current study: 9 PD, 6 PDD, and 5 DLB, as well as 10 age-matched control donors. During life, all donors provided written informed consent for the use of their brain tissue and medical records for research purposes. PD, PDD and DLB donors were included in collaboration with the Netherlands Brain Bank (NBB; http://brainbank.nl). Based on available clinical information, PDD was diagnosed if dementia developed at least a year after the onset of the motor symptoms, whereas DLB was defined if dementia developed within one year from the onset of motor symptoms^30^. As the neuropathological hallmarks of PDD and DLB are similar^1,31^, they were combined as one PDD/DLB group in the present study. The controls were included at the department of Anatomy and Neurosciences, Amsterdam UMC, following the Normal Aging Brain Collection Amsterdam (NABCA; http://nabca.eu) pipeline^32^. All donors underwent post-mortem *in situ* MRI, brain autopsy and dissection and neuropathological diagnosis by an expert neuropathologist (JMR) according to the international guidelines of the Brain Net Europe (BNE) consortium^33^. The study design is summarized in Fig 1.

**Fig 1.**
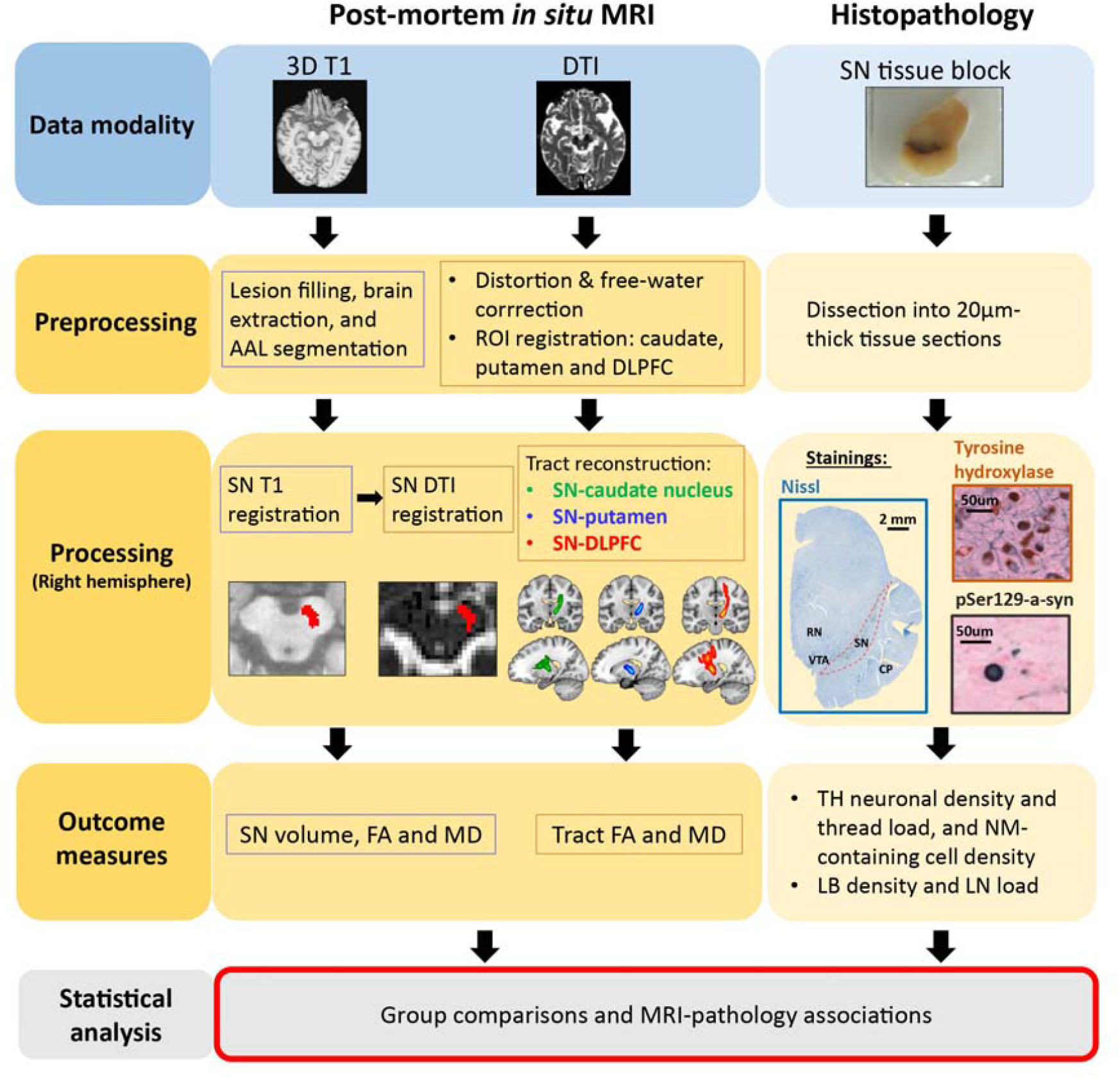
Flow chart of study design. Abbreviations: DTI, diffusion tensor imaging; SN, substantia nigra; AAL, automated anatomical labeling; DLPFC, dorsolateral prefrontal cortex; IHC, immunohistochemistry; TH, tyrosine hydroxylase; pSer129-α-syn, phosphorylated-Serine129-α-synuclein; FA, fractional anisotropy; MD, mean diffusivity; NM, neuromelanin; LB, Lewy body; LN, Lewy neurite.

### MRI acquisition

Post-mortem MRI data was acquired by whole-brain *in situ* (brain still in cranium) scanning on a whole-body 3T MR scanner (Signa-MR750, General Electric Medical Systems, United States) with an eight-channel phased-array head-coil.^32^ T1-weighted images (T1w) were acquired using a sagittal 3D T1-weighted fast spoiled gradient echo sequence (GRE) with the following parameters: repetition time (TR)/echo time (TE)/ inversion time (TI) = 7/3/450 ms, flip angle = 15°, slice thickness= 1 mm, in-plane resolution= 1.0 × 1.0 mm^2^. A sagittal 3D fluid attenuation inversion recovery (FLAIR) was acquired with TR/TE/TI= 8000/130/2000-2250 ms, slice thickness 1.2 mm, in-plane resolution= 1.11 × 1.11 mm^2^. In addition, the inversion time of the FLAIR sequence was optimized per case to account for variable CSF suppression due to post-mortem delay (PMD). Diffusion tensor imaging (DTI) was acquired by axial 2D echo-planar imaging with diffusion gradients applied in 30 non-collinear directions, TR/TE= 7400/92 ms, slice thickness 2.0 mm, in-plane resolution=2.0 × 2.0 mm^2^, and b=1000 s/mm^2^, and 5 b0 images. To allow for geometric distortion correction, b0 images with reversed phase-encode direction were obtained as well.

### MRI analysis

#### Structural image processing and SN atlas registration

To minimize the impact of age-related white matter abnormalities (e.g., vascular change) on automated segmentations, the 3D T1 images were lesion-filled,^34^ as previously described.^35^ Subsequently, normalized brain volumes of the whole brain, white matter, and gray matter were estimated from 3D T1 images using SIENAX,^36^ FMRIB Software Library (FSL) tools version 6.0.4 (https://fsl.fmrib.ox.ac.uk/fsl/). The SN pars compacta (SNpc) and reticulate (SNpr) were defined in 3D T1 space using an existing probabilistic atlas of the human subcortical brain nuclei^37^, as illustrated in Fig 1, and later on combined as a whole SN to enable sufficient voxels for tractography in diffusion MRI analysis. In addition, each hemisphere was parcellated into 39 anatomical regions using the automated anatomical labeling (AAL) atlas, whereas 14 subcortical regions were delineated from deep gray matter and parcellated using model-based segmentation/registration tool (FIRST)^38^. The caudate nucleus, putamen and DLPFC, along with the SN, were selected and transformed from 3D T1 to diffusion space using Boundary-Based registration, linear and non-linear registration (FSL) for further diffusion analysis^39^.

#### DTI pre-processing, free water correction and probabilistic tractography

DTI was first corrected for eddy current induced geometric distortion and fitted for bi-tensors diffusion maps following by free-water correction^40–42^, which has shown to yield a more accurate tensor estimation in the voxels by attenuating the partial volume effect^43,44^. The bi-tensor model for free water correction was performed using an open resource, python-based script of DIPY^40,45,46^, deriving free water corrected FA and MD maps. Subsequently, diffusion orientation distributions were modeled using FDT (part of FSL 6.0.4). The SN tracts were determined using probabilistic tractography. For this, BEDPOSTX, a function from FDT, was used to model the distribution of fiber orientations at each voxel yielding the voxel-wise diffusion orientations for performing probabilistic tractography. Probabilistic tractography was performed using ProbTrackX2 (FDT, FSL 6.0.4) with default settings and 5000 sampling fibers. As tissue blocks and subsequent histopathological outcome measures were only available in the right hemisphere, the tractography was performed only in the right hemisphere. The SN tracts were reconstructed with the SN as seed region of interest (ROI) and the caudate nucleus, putamen and DLPFC as target ROIs, resulting in three tracts: SN-caudate nucleus, SN-putamen, and SN-DLPFC. All tracts were thresholded with 0.2% to remove spurious tracts and retain anatomically correct tracts. After this, the tracts were binarized, in line with previous work^47^, and overlaid on free water-corrected diffusion maps to obtain the FA and MD of the tracts and of the SN (Fig 1).

### SN tissue sampling

After post-mortem *in situ* MRI, the donors were transported to the mortuary for craniotomy. The left hemisphere was instantly dissected and preserved for molecular and biochemistry analysis, whereas the right hemisphere was fixed in 4% formalin for four weeks and subsequently dissected for paraffin embedding and (immuno)histochemical analysis.^32,48^ Based on the BNE sampling protocol, the right SN block was cut at the level of the midbrain. The block was subsequently paraffin embedded, followed by immunohistochemistry.

#### Immunohistochemistry for detection of dopaminergic neurons and pSer129 α-Synuclein pathology

For detailed methods, see Supplementary Materials. In brief, paraffin-embedded tissue blocks of the SN were cut into 3x 20 µm-thick consecutive sections and stained for mouse-anti tyrosine hydroxylase (TH, dilution 1:800, Immunostar, Hudson, USA), or rabbit-anti phosphorylated-Ser129 α-synuclein (pSer129-asyn, clone EP1536Y, dilution 1:4000, Abcam, Cambridge, UK), and Nissl bodies with Thionin (Sigma-Aldrich, Missouri, United States). The primary antibodies were incubated at 4°C overnight. Finally, both TH and pSer129-α-syn were visualized with Vector SG grey (Vector, California, United State) followed by counter-staining with fast red (Vector, California, United State), and dehydrated in a series of ethanol, xylene, then mounted with Entellan.

#### SN delineation, and quantification of dopaminergic neurons and α-Syn pathology

Immunostained sections were digitally scanned with the Vectra Polaris Quantitative Pathology Imaging System (PerkinElmer, USA) using a 200x magnification. SN delineation and quantification of TH+ and neuromelaning-contraining neurons, TH+ fibers and a-syn pathology were performed using Qupath Software Version 0.2.3^49^. Based on previous literature^50–53^, the SN was annotated on Nissl images based on neighboring anatomical landmarks to avoid disease-related dopaminergic loss affecting the selected area. In summary, the ventral tegmental area, which are densely clustered, smaller, and lesser stained cell groups, served as the medial border; the red nucleus and white matter tracts served as the dorsal border of the SN; the cerebral peduncle was used as the ventral outline, which extends to the lateral side of the midbrain (Supplementary Fig 1). The SN delineation was transferred to consecutively stained sections of TH and pSer129-α-syn, after which, TH and pSer129-α-syn immunoreactivity, as well as neuromelanin-positive neurons, were quantified using in-house QuPath scripts. Detailed description of the scripts can be found in Supplementary Methods and Fig 1. To summarize, the pathological outcome measures included TH positive (TH+) neuronal density (count/mm^2^), TH+ thread load (% area), neuromelanin-containing cell density (count/mm^2^), LB density (count/mm^2^), and LN load (% area).

### Statistical analysis

Statistical analysis was performed using IBM SPSS 22.0 for Windows (SPSS, Inc., Chicago, IL, USA). All statistical variables were tested for normality. Chi-square test was used to investigate group difference between controls, PD and PDD/DLB donors for categorical variables. General linear models (GLM) were used for the aforementioned group differences in MRI-derived outcome measures (SN volume, FA and MD, as well as FA and MD of the specific tracts), and histopathological outcome measures. We applied linear regression models to examine the associations between the above-mentioned MRI and histopathology-derived outcome measures. Age, gender and PMD were included as covariates in all analysis. The regression models performed in two ways: 1) within the whole cohort, to address the pathological sensitivity of identified MRI measures irrespective of disease status; 2) only within the PD and PDD/DLB group. Disease duration was included in statistical analysis that involved the PD and PDD/DLB groups^54^. Group comparisons and correlation analysis of the whole cohort were followed by false discovery rate (FDR) correction for multiple comparisons^55^, while the correlation analysis within the PD+PDD/DLB group were not corrected due to the smaller sample size and exploratory nature. The graphical illustrations were made with Biorender (https://biorender.com/), R (https://www.rstudio.com/) and Graphpad (https://www.graphpad.com/).

### Data sharing

The data that support the findings of this study are available upon reasonable request.

## Results

### Clinical, radiological and pathological characteristics

Group demographics are summarized in Table 1. For detailed donor information, see Supplementary Table 1. In general, no differences in gender or PMD were found between groups. PD and PDD/DLB donors were older than controls, but no age difference between PD and PDD/DLB was observed. For radiological characteristics, no differences were found in normalized whole brain, white and grey matter volume after correction for age, gender and PMD. Lastly, disease duration was shorter in PDD/DLB compared to PD donors (*p<* .01), due to the shorter disease duration of DLB donors (14 ± 5 years in PDD v.s. 5± 2 years in DLB).

**Table 1.** Group demographics.

|  | Controls | PD+PDD/DLB | PD | PDD/DLB |
| --- | --- | --- | --- | --- |
| **Clinical characteristics** |  |  |  |  |
| N | 10 | 20 | 9 | 11 |
| Gender F/M (%M) | 5/5 (50%) | 8/12 (60%) | 3/6 (67%) | 5/6 (55%) |
| Age at death<br>years, mean ± SD | $70 \pm 7$ | $80 \pm 8^{**}$ | $80 \pm 9^*$ | $80 \pm 8^*$ |
| Disease duration<br>years, mean ± SD | - | $13,3 \pm 6,8$ | $17,2 \pm 4,4$ | $9,8 \pm 7^{\#}$ |
| PMD<br>mean (hr:min) ± SD<br>(hr:min) | $8:30 \pm 1:23$ | $7:43 \pm 2:02$ | $7:54 \pm 2:27$ | $7:35 \pm 1:43$ |
| **Radiological characteristics** |  |  |  |  |
| N | 10 | 20 | 9 | 11 |
| NWMV (ml) | $753,4 \pm 47,7$ | $705,1 \pm 41,0$ | $715,2 \pm 53,2$ | $683,1 \pm 15,1$ |
| NGMV (ml) | $716,9 \pm 39,8$ | $697,6 \pm 50,3$ | $716,9 \pm 44,7$ | $695,5 \pm 37,0$ |
| NBV (ml) | $1470,2 \pm 69,9$ | $1402,7 \pm 79,4$ | $1432,1 \pm 85,0$ | $1378,6 \pm 69,1$ |
| **Pathological characteristics** |  |  |  |  |
| **Braak LB stage N** | 10 | 20 | 9 | 11 |
| 0/1/2/3/4/5/6 | 10/0/0/0/0/0/0 | 0/0/0/0/1/1/18 | 0/0/0/0/1/1/7 | 0/0/0/0/0/0/11 |
| **Braak NFT stage N** | 10 | 20 | 9 | 11 |
| 0/1/2/3/4/5/6 | 2/5/3/0/0/0/0 | 0/2/9/3/6/0/0 | 0/2/5/1/1/0/0 | 0/0/4/2/5/0/0 |
| **Thal phase N** | 10 | 20 | 9 | 11 |
| 0/1/2/3/4/5 | 2/4/3/1/0/0 | 1/4/4/7/2/2 | 1/3/3/2/0/0 | 0/1/1/5/3/1 |
| **ABC score N** | 10 | 20 | 9 | 11 |
| A 0/1/2/3 | 2/7/1/0 | 1/8/7/4 | 1/6/2/0 | 0/2/5/4 |
| B 0/1/2/3 | 2/8/0/0 | 0/11/9/0 | 0/7/2/0 | 0/4/7/0 |
| C 0/1/2/3 | 10/0/0/0 | 12/2/6/0 | 8/1/0/0 | 4/1/6/0 |

Abbreviations: PD, Parkinson's disease; PDD, Parkinson's disease with dementia; DLB, dementia with Lewy bodies; F, female; M, male; SD, standard deviation; PMD, post-mortem delay; hr, hour; min, minutes; NWMV, normalized white matter volume; NGMV, normalized grey matter volume; NBV, normalized brain volume; ml, milliliter; LB, Lewy body; NFT, neurofibrillary tangle; ABC, amyloid, Braak, CERAD (Consortium to Establish a Registry for Alzheimer's Disease). \* $p < .05$ compared to controls, \*\* $p < .001$ compared to controls, # $p < .05$ compared to PD.

### Microstructural integrity of the SN and its tracts

Group comparisons of MRI outcome measures are illustrated in Fig 3. Compared to controls, PD+PDD/DLB combined showed increased MD of the SN (*p=*.048), but no difference in SN FA or volume was found (respectively, *p*=.306 and *p*=.344) (Fig 2A). There were no group differences between PD and controls, PDD/DLB and controls, or PD and PDD/DLB in SN volume, FA and MD (all p>0.05). (Supplementary Table 2)

**Fig 2.**
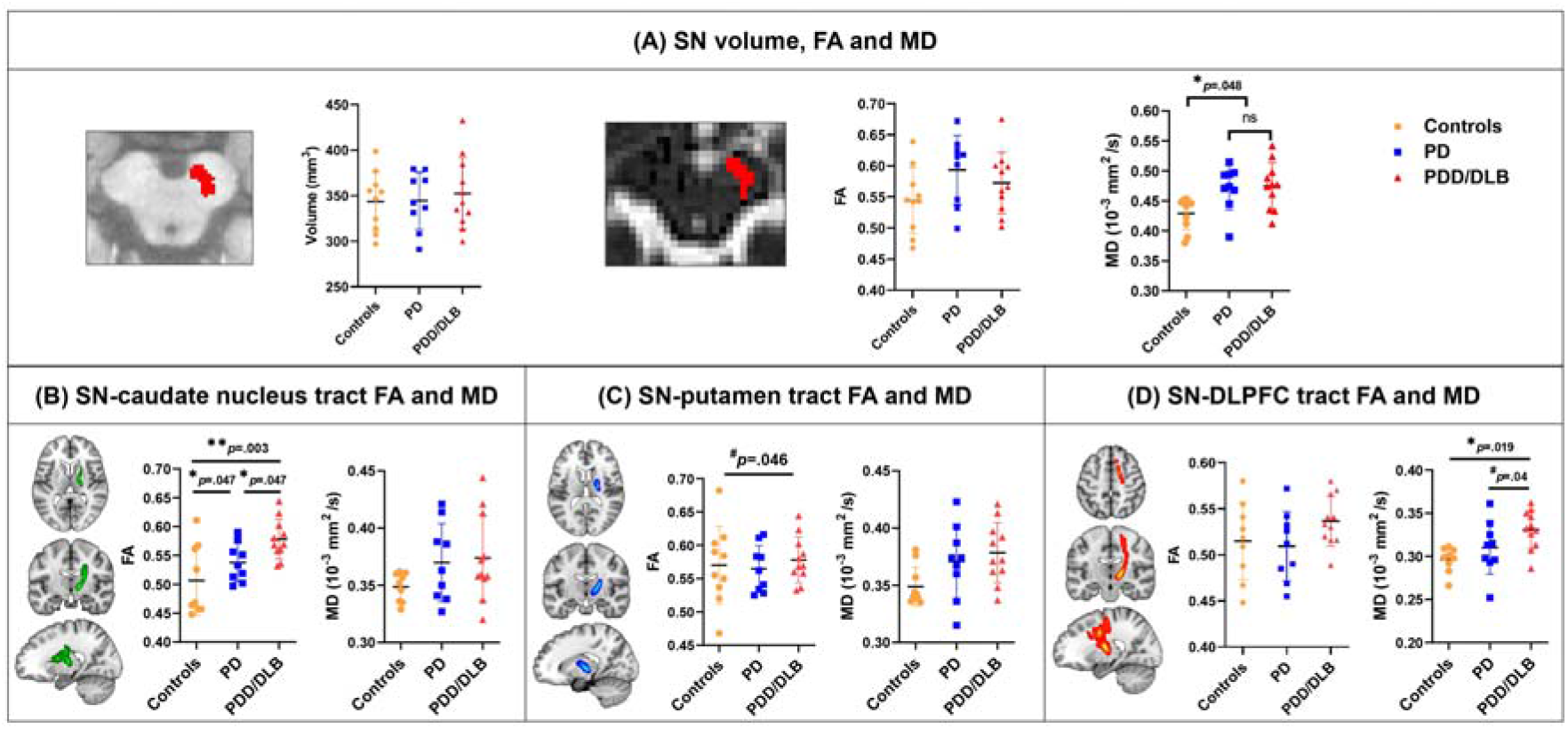
Group comparisons in MRI measures within the SN (A) and tracts of SN-caudate nucleus (B), SN-putamen (C) and SN-DLPFC (D). Abbreviations: PD, Parkinson’s disease; PDD, Parkinson’s disease with dementia; DLB, dementia with Lewy bodies; SN, substantia nigra; FA, fractional anisotropy; MD, mean diffusivity. \**p<*0.05 corrected, \*\**p<*0.01 corrected, #*p<*.05 uncorrected. ns=not significant.

**Fig 3.**
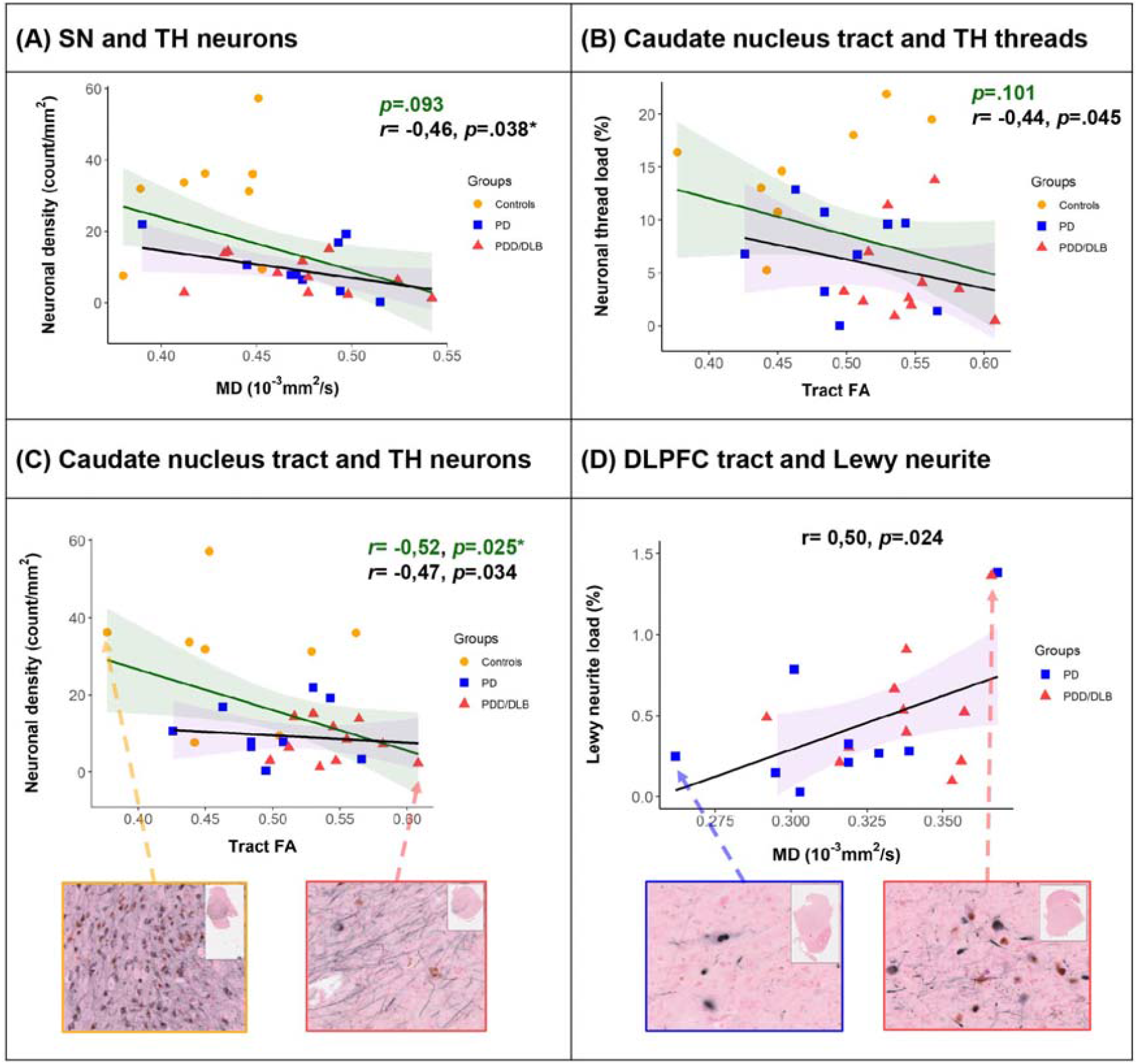
MRI-histopathology associations within the SN (A), and between tract measures and nigral pathology (B)(C)(D). Abbreviations: PD, Parkinson’s disease; PDD, Parkinson’s disease with dementia; DLB, dementia with Lewy bodies; SN, substantia nigra; TH, tyrosine hydroxylase; MD, mean diffusivity; FA, fractional anisotropy. \**p<*.05, corrected.

For the microstructural integrity of the tracts, compared to controls, PD+PDD/DLB showed an increased FA in SN-caudate nucleus tract (*p=*.012), and a trend of increased FA in the tracts between the SN and putamen (*p=*.057), while no differences were found for MD (*p=*.557 for SN-caudate, and *p=*.119 for SN-putamen). In addition, PD+PDD/DLB showed an increased MD in the SN-DLPFC tract (*p=*.047), which did not survive correction for multiple comparison. No difference in FA in the SN-DLPFC tract was found (*p=*.575).

PDD/DLB exhibited higher FA in the SN-caudate nucleus tract compared to controls (*p=*.003) (Fig 2B), but the SN-putamen tract (*p=*.046) did not survive correction for multiple comparisons (Fig 2C). In addition, MD of the SN-DLPFC tract was higher in PDD/DLB than controls (*p=*.019) (Fig 2D). Compared to PD, PDD/DLB showed higher FA of the SN-caudate nucleus tract (*p=*.047), and higher MD of the SN-DLPFC tract (*p=*.04). No differences were found in the SN-putamen tract for PDD/DLB compared to PD (*p=*.750). When comparing PD and controls, only the SN-caudate nucleus tract showed increased FA in PD (*p=*.047) (Fig 2C). In summary, among the three groups, PDD/DLB showed more severely altered microstructure within the SN-caudate nucleus tract, as well as within the SN-DLPFC tract, but not within the SN-putamen tract.

### Histopathological features in the SN of PD and PDD/DLB donors

No significant differences were found between PD and PDD/DLB in both SN LB density and LN load. (Supplementary Fig 2A-B). Compared to controls, PD donors showed 66% less TH+ neurons (*P=*.014) and 55% less TH+ thread load (*p=*.049), as well as 82% less neuromelanin-containing neurons (*p=*.002). In turn, PDD/DLB showed 74% less TH+ neurons (*p=*.006) and 69% less TH+ threads (*p=*.012), as well as 83% less neuromelanin-containing neurons (*p=*.002) compared to controls. No differences were found between PD and PDD/DLB. (Supplementary Fig 2C-E)

### Pathological correlates of microstructural alterations in the SN and its tracts

To identify the pathological correlate(s) that underlie the above mentioned MRI-derived findings, we included the SN and the tracts that showed significant group differences between PD, PDD/DLB and controls, namely MD of the SN, FA of the SN-caudate nucleus tract, and MD of the SN-DLPFC tract.

Within the SN, increased MD correlated with both reduced TH+ and neuromelainin-containing neuronal density in PD+PDD/DLB (*r=* -.46, *p=* .038 and *r=* -.51, *p=* .022, respectively), but not in the whole cohort (*p=* .093 and *p=*,327, respectively) (Fig 3A). Increased FA of the SN-caudate nucleus correlated with reduced TH+ thread load in PD+PDD/DLB (*r=* -0.44, *p=* .045), but not the whole cohort (*p=* .101) (Fig 3B); and with reduced TH+ neuronal density in both PD+PDD/DLB (*r=* -0.47, *p=* .034) and the whole cohort (*r=* -0.52, *p=* .025), though the correlation in PD+PDD/DLB did not survive the multiple comparison (Fig. 3C). In addition, increased MD of the SN-DLPFC correlated with increased LN load (*r=* .50, *p=* .024) in the PD+PDD/DLB group (Fig 3D). Supplementary Table 3 summarized the Pearson’s r and p-values of the correlations between MRI and SN histopathological outcome measures.

In summary dopaminergic cell loss is associated with an increased FA of the SN-caudate nucleus tract, and an increased LN load is associated with an increased MD of the SN-DLPFC tract.

## Discussion

This is the first study to address the underlying pathological substrates of MRI-measured alterations of the SN and its tracts to the dorsal striatum and DLPFC in PD and PDD/DLB. We showed that SN MD and SN-caudate nucleus tract FA were increased in PD and PDD/DLB, and SN-DLPFC tract MD was increased only in PDD/DLB. Furthermore, microstructural alterations of the SN and SN-caudate nucleus tract was related to decreased SN dopaminergic neuronal density, and alteration of the SN-DLPFC tract was related to increased SN LN load.

In line with the literature, we found reduced microstructural integrity of the SN in PD+PDD/DLB compared to controls^16,56^. In previous studies, increased SN MD was shown to be a promising biomarker in following disease progression, and was linked to bradykinesia, cognitive status and dopaminergic deficits^19,21,43,56^. Here we show that SN MD is driven by reduced dopaminergic neuronal density in the SN, providing the missing link between clinical/radiological and histopathological studies. Presumably, the loss of dopaminergic neurons enlarges the extracellular space, thus increasing water diffusivity. As such, SN MD is a potential imaging marker for dopaminergic neuronal loss within this region, although further investigation in a larger cohort is warranted. Previous studies also discovered progressive SN volume loss in PD^57^. However, we did not find SN volume loss in PD or PDD/DLB in our cohort. This may be due to the fact that the SNpc and SNpr were combined in the current study to enable sufficient voxels for the tractography. As dopaminergic loss is more pronounced in the SNpc, which contributes to the volume loss on MRI, combining both regions may have diluted this observation.

Microstructural alterations in PD and PDD/DLB were found in SN-caudate nucleus and SN-DLPFC tracts. We did not find an alteration in the SN-putamen tract in PD or PDD/DLB, which may be surprising since the posterior putamen has been shown to be more and earlier affected than the caudate nucleus^58–61^. However, this may be due to the fact that the PD and PDD/DLB donors in our study were late-to-end stage patients, and dopaminergic loss in the putamen may has reached a ceiling effect, providing limited variability for an association with microstructural integrity.

Rather than an increase in FA of the SN-caudate nucleus tract, as shown in our study, some previous *in-vivo* studies found a decreased FA of the nigrostriatal tract^21,62–64^. In these studies, PD patients were sampled at early to middle stages of their disease (Hoehn and Yahr stage of 1-3) with an average age of 70, and disease duration of 3-10 years^16,65^. In the current study, PD and PDD donors were at late to end stages of their disease, with an average age of 80 years and a relatively long disease duration of 8-23 years. The sampled disease stages may influence the observation of FA, as higher age and longer disease duration (as in our study with end-stage brain donors) are associated with reduced axonal density^60,66,67^. An overall axonal loss in the white matter, may result in a reduced number of crossing fibers and in turn coherently elongated principal diffusion axis and an increased FA^67,68^. An alternative explanation, or concomitant phenomena, is that the nigrostriatal neurons in PD undergo an inflammatory process prior to neuronal death^69^, which leads to intracellular water influx and decreased extracellular water^70^. This results in decreased diffusivity perpendicular to the axon, secondary to more compacted axons, and in turn an increased anisotropy^68,71^. The higher FA of the SN-caudate nucleus tract in our study may thus reflect an additional axonal degenerative process.

An increased MD of the SN-DLPFC tract in PDD/DLB, and not in PD, indicates the involvement of this tract in dementia-related symptoms, and may have the potential as an imaging marker in disease progression, particularly in the transition from PD to PDD. Previous studies in PD showed widespread microstructural damage in frontal white matter, which associated with executive and visuospatial disability, as well as with axial disturbances, namely postural instability and gait disturbances^72,73^. Our study shows that increased MD of the SN-DLPFC tract is associated with α-syn accumulation within the SN, rather than dopaminergic loss. In support of this, cognitive deficits and axial disturbances are ascribed to non-dopaminergic functions, as they are not responsive to Levodopa treatment74,75,76,77.

Despite the novel approach of the current study with a combined MRI and histopathology method, some limitations should be addressed. Although a sample size of 30 cases is considered large for *in situ* MRI-pathology studies, our results require replication in a larger cohort to further validate the MRI-pathological associations. Aside from dopaminergic fibers, the TH threads may also include the noradrenergic fibers that innervate the SN from the locus coeruleus (LC), as TH is the precursor of both dopamine and noradrenaline^78–81^. A reduction in LC noradrenergic activity may also result in reduced activity of SN neurons^82–84^. In addition, PDD and DLB often show Alzheimer’s disease (AD) co-pathology, such as amyloid-beta (Aβ) plaques and phosphorylated-tau (p-tau) neurofibrillary tangles (NFT)^85,86^, and the influence of these markers should be taken into consideration. As the pars compacta and reticulata disproportionally contribute to neuronal loss in the SN^87^, combining these regions could have led to a lack of measurable SN volume loss on MRI. Future studies are advised to leverage a higher resolution DTI and/or neuromelain-sensitive imaging to specifically assess the dopaminergic alterations within the SN pars compacta.

In conclusion, our study shows that diffusion MRI markers are able to capture microstructural alterations of the SN and its tracts in PD, and more pronounced in PDD/DLB. More importantly, these MRI-measured alterations were associated with nigral pathology, specifically dopaminergic loss and LN load, as shown in Fig 4. Our findings provide fundamental knowledge on the pathological sensitivity of MRI microstructure, which refines the interpretation of neuroimaging results. These may aid the development of reliable diagnostic/monitoring indicators for cognitive state in PD and dementia in PDD/DLB.

**Fig 4.**
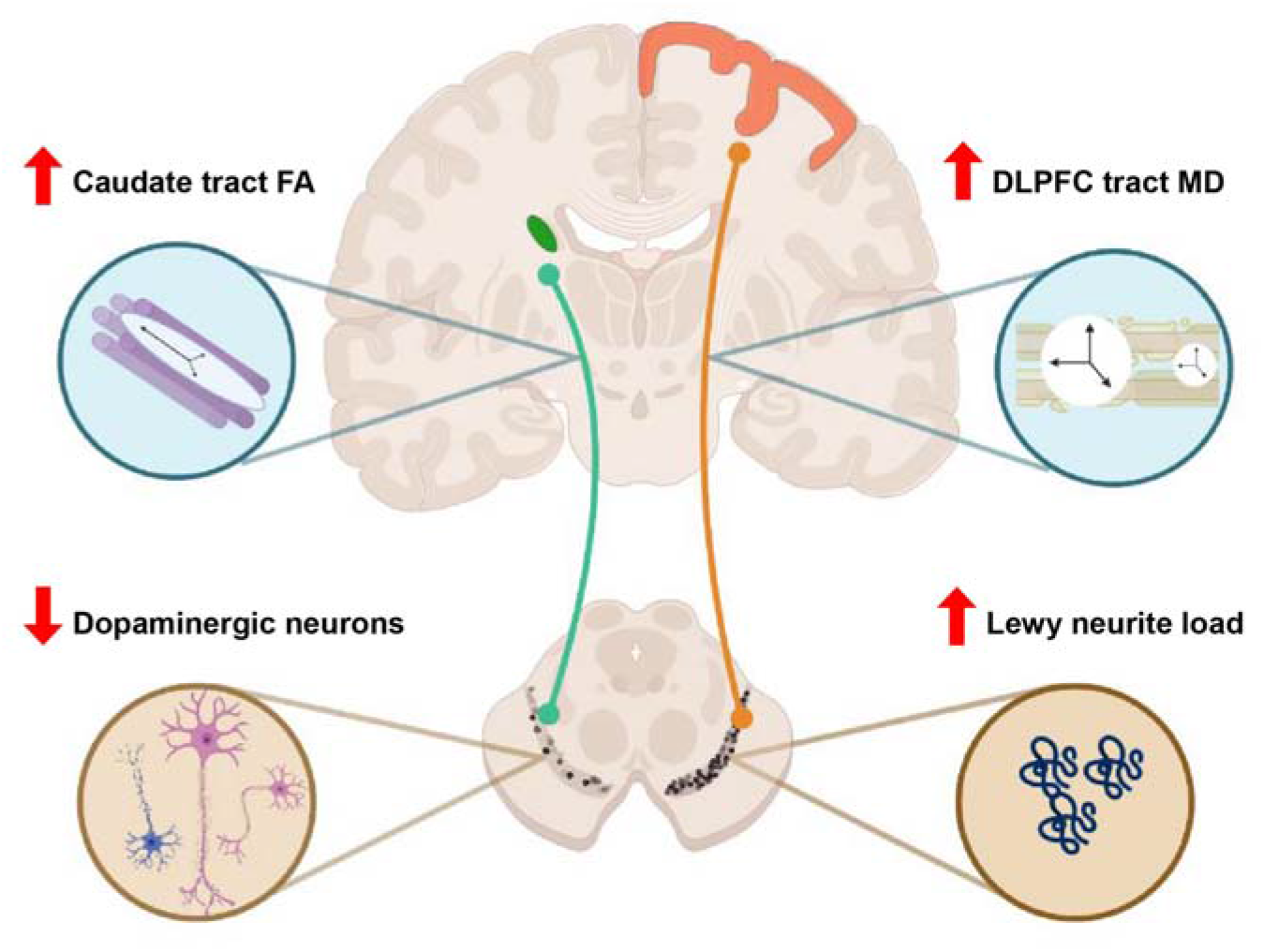
Summary model of the pathological sensitivity of SN projections. Pathological change within the substantia nigra (SN), specifically dopaminergic cell loss and increased Lewy neurite (LN) load are associated with MRI-DTI microstructural alteration. Dopaminergic cell loss is associated with an increased FA of tracts between the SN and caudate nucleus, and an increased LN load is associated with an increased MD of tracts between the SN and DLPFC.

## Acknowledgements

We would like to thank all brain donors and their caregivers for donating their brain to scientific research, as well as the Netherlands Brain Bank (www.brainbank.nl and the Normal Aging Brain Collection Amsterdam (www.nabca.eu; NABCA) MRI and autopsy teams. Special thanks to Angela Ingrassia and Allert Jonker for helping in the lab with the immunohistochemical stainings.

## Author’s role

(1) Research project: A. Conception, B. Organization, C. Execution; (2) Analysis: A. Design, B. Execution, C. Review and Critique; (3) Manuscript Preparation: A. Writing of the First Draft, B. Review and Critique

Chen-Pei Lin: (1)ABC, (2)ABC, (3) A

Lydian EJ Knoop: (1)BC, (2)AB

B Irene Frigerio: (2)A, (3)B

John GJM Bol: (1)C, (3)B

Annemieke JM Rozemuller: (1)C

Henk W Berendse: (1)A, (3)B

Petra JW Pouwels: (1)A, (2)C, (3)B

Wilma DJ van de Berg: (1)AB, (2)AC,

(3)B Laura E Jonkman: (1)AB, (2)AC, (3)AB

## References

1. Tolosa E, Wenning G, Poewe W. The diagnosis of Parkinson’s disease. Lancet Neurol. 2006;5(1). DOI:10.1016/S1474-4422(05)70285-4

2. Aarsland D, Batzu L, Halliday GM, et al. Parkinson disease-associated cognitive impairment. Nat Rev Dis Prim. 2021;7(1). DOI:10.1038/s41572-021-00280-3

3. Sawamoto N, Piccini P, Hotton G, Pavese N, Thielemans K, Brooks DJ. Cognitive deficits and striato-frontal dopamine release in Parkinson’s disease. Brain. 2008;131(5). DOI:10.1093/brain/awn054

4. Rinne JO, Portin R, Ruottinen H, et al. Cognitive impairment and the brain dopaminergic system in Parkinson disease: [18F]fluorodopa positron emission tomographic study. Arch Neurol. 2000;57(4). DOI:10.1001/archneur.57.4.470

5. Severiano e Sousa C, Alarcão J, Pavão Martins I, Ferreira JJ. Frequency of dementia in Parkinson’s disease: A systematic review and meta-analysis. J Neurol Sci. 2022;432. DOI:10.1016/j.jns.2021.120077

6. Aarsland D, Zaccai J, Brayne C. A systematic review of prevalence studies of dementia in Parkinson’s disease. Mov Disord. 2005;20(10). DOI:10.1002/mds.20527

7. Jellinger KA. Morphological differences between dementia with Lewy bodies and Parkinson’s disease-dementia. Neuropathol Appl Neurobiol. 2022;48(2). DOI:10.1111/nan.12708

8. Tolosa E, Garrido A, Scholz SW, Poewe W. Challenges in the diagnosis of Parkinson’s disease. Lancet Neurol. 2021;20(5). DOI:10.1016/S1474-4422(21)00030-2

9. Li T, Le W. Biomarkers for Parkinson’s Disease: How Good Are They? Neurosci Bull. 2020;36(2). DOI:10.1007/s12264-019-00433-1

10. Lotankar S, Prabhavalkar KS, Bhatt LK. Biomarkers for Parkinson’s Disease: Recent Advancement. Neurosci Bull. 2017;33(5). DOI:10.1007/s12264-017-0183-5

11. Wang J, Zhang F, Zhao C, et al. Investigation of local white matter abnormality in Parkinson’s disease by using an automatic fiber tract parcellation. Behav Brain Res. 2020;394. DOI:10.1016/j.bbr.2020.112805

12. Vaillancourt DE, Prodoehl J, Abraham I, et al. High-resolution diffusion tensor imaging in the substantia nigra of de novo Parkinson disease. Neurology. 2009;72(16). DOI:10.1212/01.wnl.0000340982.01727.6e

13. Zhan W, Kang GA, Glass GA, et al. Regional alterations of brain microstructure in Parkinson’s disease using diffusion tensor imaging. Mov Disord. 2012;27(1). DOI:10.1002/mds.23917

14. Péran P, Cherubini A, Assogna F, et al. Magnetic resonance imaging markers of Parkinson’s disease nigrostriatal signature. Brain. 2010;133(11). DOI:10.1093/brain/awq212

15. Theisen F, Leda R, Pozorski V, et al. Evaluation of striatonigral connectivity using probabilistic tractography in Parkinson’s disease. NeuroImage Clin. 2017;16. DOI:10.1016/j.nicl.2017.09.009

16. Zhang Y, Burock MA. Diffusion Tensor Imaging in Parkinson’s Disease and Parkinsonian Syndrome: A Systematic Review. Front Neurol. 2020;11. DOI:10.3389/fneur.2020.531993

17. Wen Q, Mustafi SM, Li J, et al. White matter alterations in early-stage Alzheimer’s disease: A tract-specific study. Alzheimer’s Dement Diagnosis, Assess Dis Monit. 2019;11:576–587. DOI:10.1016/j.dadm.2019.06.003

18. Mole JP, Subramanian L, Bracht T, Morris H, Metzler-Baddeley C, Linden DEJ. Increased fractional anisotropy in the motor tracts of Parkinson’s disease suggests compensatory neuroplasticity or selective neurodegeneration. Eur Radiol. 2016;26(10). DOI:10.1007/s00330-015-4178-1

19. Loane C, Politis M, Kefalopoulou Z, et al. Aberrant nigral diffusion in Parkinson’s disease: A longitudinal diffusion tensor imaging study. Mov Disord. 2016;31(7). DOI:10.1002/mds.26606

20. Shang S, Li D, Tian Y, et al. Hybrid PET-MRI for early detection of dopaminergic dysfunction and microstructural degradation involved in Parkinson’s disease. Commun Biol. 2021;4(1). DOI:10.1038/s42003-021-02705-x

21. Zhang Y, Wu IW, Buckley S, et al. Diffusion tensor imaging of the nigrostriatal fibers in Parkinson’s disease. Mov Disord. 2015;30(9). DOI:10.1002/mds.26251

22. Foltynie T, Brayne CEG, Robbins TW, Barker RA. The cognitive ability of an incident cohort of Parkinson’s patients in the UK. The CamPaIGN study. Brain. 2004;127(3). DOI:10.1093/brain/awh067

23. Janvin CC, Aarsland D, Larsen JP. Cognitive predictors of dementia in Parkinson’s disease: A community-based, 4-year longitudinal study. J Geriatr Psychiatry Neurol. 2005;18(3). DOI:10.1177/0891988705277540

24. Williams-Gray CH, Mason SL, Evans JR, et al. The CamPaIGN study of Parkinson’s disease: 10-year outlook in an incident population-based cohort. J Neurol Neurosurg Psychiatry. 2013;84(11). DOI:10.1136/jnnp-2013-305277

25. Mahieux F, Fénelon G, Flahault A, Manifacier MJ, Michelet D, Boller F. Neuropsychological prediction of dementia in Parkinson’s disease. J Neurol Neurosurg Psychiatry. 1998;64(2). DOI:10.1136/jnnp.64.2.178

26. Okubo Y, Suhara T, Suzuki K, et al. Decreased prefrontal dopamine D1 receptors in schizophrenia revealed by PET. Nature. 1997;385(6617). DOI:10.1038/385634a0

27. Abi-Dargham A, Mawlawi O, Lombardo I, et al. Prefrontal Dopamine D1 Receptors and Working Memory in Schizophrenia. J Neurosci. 2002;22(9). DOI:10.1523/jneurosci.22-09-03708.2002

28. Rakshi JS, Uema T, Ito K, et al. Frontal, midbrain and striatal dopaminergic function in early and advanced Parkinson’s disease. A 3D [18F]dopa-PET study. Brain. 1999;122(9). DOI:10.1093/brain/122.9.1637

29. Kaasinen V, Någren K, Hietala J, et al. Extrastriatal dopamine D2 and D3 receptors in early and advanced Parkinson’s disease. Neurology. 2000;54(7). DOI:10.1212/WNL.54.7.1482

30. McKeith IG. Clinical Diagnostic Criteria for Dementia with Lewy Bodies. In: Dementia with Lewy Bodies. ; 2017. DOI:10.1007/978-4-431-55948-1_5

31. Emre M, Aarsland D, Brown R, et al. Clinical diagnostic criteria for dementia associated with Parkinson’s disease. Mov Disord. 2007;22(12). DOI:10.1002/mds.21507

32. Jonkman LE, Graaf YG de, Bulk M, et al. Normal Aging Brain Collection Amsterdam (NABCA): A comprehensive collection of postmortem high-field imaging, neuropathological and morphometric datasets of non-neurological controls. NeuroImage Clin. 2019;22. DOI:10.1016/j.nicl.2019.101698

33. Alafuzoff I, Ince PG, Arzberger T, et al. Staging/typing of Lewy body related α-synuclein pathology: A study of the BrainNet Europe Consortium. Acta Neuropathol. 2009;117(6). DOI:10.1007/s00401-009-0523-2

34. Steenwijk MD, Pouwels PJW, Daams M, et al. Accurate white matter lesion segmentation by k nearest neighbor classification with tissue type priors (kNN-TTPs). NeuroImage Clin. 2013;3. DOI:10.1016/j.nicl.2013.10.003

35. Jonkman LE, Steenwijk MD, Boesen N, et al. Relationship between β-amyloid and structural network topology in decedents without dementia. Neurology. 2020;95(5). DOI:10.1212/WNL.0000000000009910

36. Smith SM, Jenkinson M, Woolrich MW, et al. Advances in functional and structural MR image analysis and implementation as FSL. In: NeuroImage. Vol 23.; 2004. DOI:10.1016/j.neuroimage.2004.07.051

37. Pauli WM, Nili AN, Michael Tyszka J. Data Descriptor: A high-resolution probabilistic in vivo atlas of human subcortical brain nuclei. Sci Data. 2018;5. DOI:10.1038/sdata.2018.63

38. Tzourio-Mazoyer N, Landeau B, Papathanassiou D, et al. Automated anatomical labeling of activations in SPM using a macroscopic anatomical parcellation of the MNI MRI single-subject brain. Neuroimage. 2002;15(1). DOI:10.1006/nimg.2001.0978

39. Greve DN, Fischl B. Accurate and robust brain image alignment using boundary-based registration. Neuroimage. 2009;48(1). DOI:10.1016/j.neuroimage.2009.06.060

40. Pasternak O, Sochen N, Gur Y, Intrator N, Assaf Y. Free water elimination and mapping from diffusion MRI. Magn Reson Med. 2009;62(3). DOI:10.1002/mrm.22055

41. Haselgrove JC, Moore JR. Correction for distortion of echo-planar images used to calculate the apparent diffusion coefficient. Magn Reson Med. 1996;36(6). DOI:10.1002/mrm.1910360620

42. Basser PJ, Mattiello J, LeBihan D. MR diffusion tensor spectroscopy and imaging. Biophys J. 1994;66(1). DOI:10.1016/S0006-3495(94)80775-1

43. Ofori E, Krismer F, Burciu RG, et al. Free water improves detection of changes in the substantia nigra in parkinsonism: A multisite study. Mov Disord. 2017;32(10). DOI:10.1002/mds.27100

44. Bergamino M, Walsh RR, Stokes AM. Free-water diffusion tensor imaging improves the accuracy and sensitivity of white matter analysis in Alzheimer’s disease. Sci Rep. 2021;11(1). DOI:10.1038/s41598-021-86505-7

45. Garyfallidis E, Brett M, Amirbekian B, et al. Dipy, a library for the analysis of diffusion MRI data. Front Neuroinform. 2014;8(FEB). DOI:10.3389/fninf.2014.00008

46. Hoy AR, Koay CG, Kecskemeti SR, Alexander AL. Optimization of a free water elimination two-compartment model for diffusion tensor imaging. Neuroimage. 2014;103. DOI:10.1016/j.neuroimage.2014.09.053

47. Hepp DH, Foncke EMJ, Berendse HW, et al. Damaged fiber tracts of the nucleus basalis of Meynert in Parkinson’s disease patients with visual hallucinations. Sci Rep. 2017;7(1). DOI:10.1038/s41598-017-10146-y

48. Klioueva NM, Rademaker MC, Dexter DT, et al. BrainNet Europe’s Code of Conduct for brain banking. J Neural Transm. 2015;122(7). DOI:10.1007/s00702-014-1353-5

49. Bankhead P, Loughrey MB, Fernández JA, et al. QuPath: Open source software for digital pathology image analysis. Sci Rep. 2017;7(1). DOI:10.1038/s41598-017-17204-5

50. Parkkinen L, O’Sullivan SS, Collins C, et al. Disentangling the Relationship between Lewy bodies and nigral neuronal loss in Parkinson’s disease. J Parkinsons Dis. 2011;1(3). DOI:10.3233/JPD-2011-11046

51. Alberico SL, Cassell MD, Narayanan NS. The vulnerable ventral tegmental area in Parkinson’s disease. Basal Ganglia. 2015;5(2-3). DOI:10.1016/j.baga.2015.06.001

52. Dijkstra AA, Voorn P, Berendse HW, Groenewegen HJ, Rozemuller AJM, van de Berg Wdj. Stage-dependent nigral neuronal loss in incidental Lewy body and parkinson’s disease. Mov Disord. 2014;29(10). DOI:10.1002/mds.25952

53. Gibb WRG, Lees AJ. Anatomy, pigmentation, ventral and dorsal subpopulations of the substantia nigra, and differential cell death in Parkinson’s disease. J Neurol Neurosurg Psychiatry. 1991;54(5). DOI:10.1136/jnnp.54.5.388

54. de Schipper LJ, van der Grond J, Marinus J, Henselmans JML, van Hilten JJ. Loss of integrity and atrophy in cingulate structural covariance networks in Parkinson’s disease. NeuroImage Clin. 2017;15. DOI:10.1016/j.nicl.2017.05.012

55. Zarkali A, McColgan P, Leyland LA, Lees AJ, Rees G, Weil RS. Fiber-specific white matter reductions in Parkinson hallucinations and visual dysfunction. Neurology. 2020;94(14). DOI:10.1212/WNL.0000000000009014

56. Atkinson-Clement C, Pinto S, Eusebio A, Coulon O. Diffusion tensor imaging in Parkinson’s disease: Review and meta-analysis. NeuroImage Clin. 2017;16. DOI:10.1016/j.nicl.2017.07.011

57. Gaurav R, Yahia-Cherif L, Pyatigorskaya N, et al. Longitudinal Changes in Neuromelanin MRI Signal in Parkinson’s Disease: A Progression Marker. Mov Disord. 2021;36(7). DOI:10.1002/mds.28531

58. Brück A, Aalto S, Rauhala E, Bergman J, Marttila R, Rinne JO. A follow-up study on 6-[18F]Fluoro-L-dopa uptake in early Parkinson’s disease shows nonlinear progressionin the putamen. Mov Disord. 2009;24(7). DOI:10.1002/mds.22484

59. Kish SJ, Shannak K, Hornykiewicz O. Uneven Pattern of Dopamine Loss in the Striatum of Patients with Idiopathic Parkinson’s Disease. N Engl J Med. 1988;318(14). DOI:10.1056/nejm198804073181402

60. Kordower JH, Olanow CW, Dodiya HB, et al. Disease duration and the integrity of the nigrostriatal system in Parkinson’s disease. Brain. 2013;136(8). DOI:10.1093/brain/awt192

61. De La Fuente-Fernández R, Schulzer M, Kuramoto L, et al. Age-specific progression of nigrostriatal dysfunction in Parkinson’s disease. Ann Neurol. 2011;69(5). DOI:10.1002/ana.22284

62. Tan WQ, Yeoh CS, Rumpel H, et al. Deterministic Tractography of the Nigrostriatal-Nigropallidal Pathway in Parkinson’s Disease. Sci Rep. 2015;5. DOI:10.1038/srep17283

63. Planetta PJ, Schulze ET, Geary EK, et al. Thalamic projection fiber integrity in de novo Parkinson Disease. Am J Neuroradiol. 2013;34(1). DOI:10.3174/ajnr.A3178

64. Schuff N, Wu IW, Buckley S, et al. Diffusion imaging of nigral alterations in early Parkinson’s disease with dopaminergic deficits. Mov Disord. 2015;30(14). DOI:10.1002/mds.26325

65. Winston GP. The physical and biological basis of quantitative parameters derived from diffusion MRI. Quant Imaging Med Surg. 2012;2(4). DOI:10.3978/j.issn.2223-4292.2012.12.05

66. Stahon KE, Bastian C, Griffith S, Kidd GJ, Brunet S, Baltan S. Age-related changes in axonal and mitochondrial ultrastructure and function in white matter. J Neurosci. 2016;36(39). DOI:10.1523/JNEUROSCI.1316-16.2016

67. Teipel SJ, Grothe MJ, Filippi M, et al. Fractional anisotropy changes in Alzheimer’s disease depend on the underlying fiber tract architecture: A multiparametric DTI study using joint independent component analysis. J Alzheimer’s Dis. 2014;41(1):69–83. DOI:10.3233/JAD-131829

68. Budde MD, Janes L, Gold E, Turtzo LC, Frank JA. The contribution of gliosis to diffusion tensor anisotropy and tractography following traumatic brain injury: Validation in the rat using Fourier analysis of stained tissue sections. Brain. 2011;134(8). DOI:10.1093/brain/awr161

69. Cicchetti F, Brownell AL, Williams K, Chen YI, Livni E, Isacson O. Neuroinflammation of the nigrostriatal pathway during progressive 6-OHDA dopamine degeneration in rats monitored by immunohistochemistry and PET imaging. Eur J Neurosci. 2002;15(6). DOI:10.1046/j.1460-9568.2002.01938.x

70. Liang D, Bhatta S, Gerzanich V, Simard JM. Cytotoxic edema: mechanisms of pathological cell swelling. Neurosurg Focus. 2007;22(5). DOI:10.3171/foc.2007.22.5.3

71. Sotak CH. The role of diffusion tensor imaging in the evaluation of ischemic brain - A review. NMR Biomed. 2002;15(7-8). DOI:10.1002/nbm.786

72. Gu Q, Huang P, Xuan M, et al. Greater Loss of White Matter Integrity in Postural Instability and Gait Difficulty Subtype of Parkinson’s Disease. Can J Neurol Sci. 2014;41(6). DOI:10.1017/cjn.2014.34

73. Gattellaro G, Minati L, Grisoli M, et al. White matter involvement in idiopathic Parkinson disease: A diffusion tensor imaging study. Am J Neuroradiol. 2009;30(6). DOI:10.3174/ajnr.A1556

74. Sethi K. Levodopa unresponsive symptoms in Parkinson disease. Mov Disord. 2008;23(SUPPL. 3). DOI:10.1002/mds.22049

75. Lang AE, Obeso JA. Challenges in Parkinson’s disease: Restoration of the nigrostriatal dopamine system is not enough. Lancet Neurol. 2004;3(5). DOI:10.1016/S1474-4422(04)00740-9

76. Alves G, Larsen JP, Emre M, Wentzel-Larsen T, Aarsland D. Changes in motor subtype and risk for incident dementia in Parkinson’s disease. Mov Disord. 2006;21(8). DOI:10.1002/mds.20897

77. Domellöf ME, Elgh E, Forsgren L. The relation between cognition and motor dysfunction in drug-naive newly diagnosed patients with Parkinson’s disease. Mov Disord. 2011;26(12). DOI:10.1002/mds.23814

78. Mejías-Aponte CA, Drouin C, Aston-Jones G. Adrenergic and noradrenergic innervation of the midbrain ventral tegmental area and retrorubral field: Prominent inputs from medullary homeostatic centers. J Neurosci. 2009;29(11). DOI:10.1523/JNEUROSCI.4632-08.2009

79. Simon H, Le Moal M, Calas A. Efferents and afferents of the ventral tegmental-A10 region studied after local injection of [3H]leucine and horseradish peroxidase. Brain Res. 1979;178(1). DOI:10.1016/0006-8993(79)90085-4

80. Jones BE, Yang T Z. The efferent projections from the reticular formation and the locus coeruleus studied by anterograde and retrograde axonal transport in the rat. J Comp Neurol. 1985;242(1). DOI:10.1002/cne.902420105

81. Szot P, Knight L, Franklin A, et al. Lesioning noradrenergic neurons of the locus coeruleus in C57Bl/6 mice with unilateral 6-hydroxydopamine injection, to assess molecular, electrophysiological and biochemical changes in noradrenergic signaling. Neuroscience. 2012;216. DOI:10.1016/j.neuroscience.2012.04.046

82. Guiard BP, El Mansari M, Merali Z, Blier P. Functional interactions between dopamine, serotonin and norepinephrine neurons: An in-vivo electrophysiological study in rats with monoaminergic lesions. Int J Neuropsychopharmacol. 2008;11(5). DOI:10.1017/S1461145707008383

83. Wang Y, Zhang QJ, Liu J, et al. Noradrenergic lesion of the locus coeruleus increases apomorphine-induced circling behavior and the firing activity of substantia nigra pars reticulata neurons in a rat model of Parkinson’s disease. Brain Res. 2010;1310. DOI:10.1016/j.brainres.2009.10.070

84. Szot P, Franklin A, Sikkema C, Wilkinson CW, Raskind MA. Sequential loss of LC noradrenergic and dopaminergic neurons results in a correlation of dopaminergic neuronal number to striatal dopamine concentration. Front Pharmacol. 2012;3 OCT. DOI:10.3389/fphar.2012.00184

85. Hepp DH, Vergoossen DLE, Huisman E, et al. Distribution and load of amyloid-b pathology in Parkinson disease and dementia with lewy bodies. J Neuropathol Exp Neurol. 2016;75(10). DOI:10.1093/jnen/nlw070

86. Lippa CF, Duda JE, Grossman M, et al. DLB and PDD boundary issues: Diagnosis, treatment, molecular pathology, and biomarkers. Neurology. 2007;68(11). DOI:10.1212/01.wnl.0000256715.13907.d3

87. Fu YH, Paxinos G, Watson C, Halliday GM. The substantia nigra and ventral tegmental dopaminergic neurons from development to degeneration. J Chem Neuroanat. 2016;76. DOI:10.1016/j.jchemneu.2016.02.001

