## Supplementary Materials for "Nigral pathology contributes to microstructural integrity of striatal and frontal tracts in Parkinson’s disease"

**Methods**

**Immunohistochemistry for Nissl bodies, tyrosine hydroxylase (TH) and phosphorylated-alpha-synuclein (pSer129- αsyn)**

Paraffin-embedded tissue blocks of the SN were cut at 20 µm (Leica Microtome) for 3 serial sections. All sections were mounted onto superfrost plus glass slides (Thermo Scientific, USA) for subsequent histological staining for Nissl bodies, and immunohistochemistry for TH and pSer129-αSyn (see main text for detailed information on antibodies).

Sections were deparaffinized and dehydrated in xylene and a graded series of ethanol. Sections for Nissl bodies were stained with Thionin directly afterwards and mounted with Entellan. Sections for TH and pSer129-αSyn subsequently underwent antigen retrieval in citrate buffer (pH 6.0) at a temperature of 95°C. Sections were first blocked for endogenous peroxidase by immersing the sections in 1% of H_2_O_2_ ,in tris-buffered saline (TBS, pH 7.4). Subsequently the sections were blocked with 0.1% Triton and 3% normal goat serum in TBS + 0.1% Triton x-100. SN sections were incubated with primary antibodies, TH or pSer129-**α**Syn. The incubated primary antibodies were diluted in 3% normal goat serum in TBS + 0.1% Triton x-100 for two nights at 4°C, followed by Immpress-HRP anti mouse or anti rabbit (Vector, California, United State) detection. Finally, both TH and αSyn were visualized with Vector SG into blue-grey (Vector, California, United State) and counterstained with nuclear fast red (Vector, California, United State), and mounting with Entellan.

**Quantification of TH and pSer129-α-Syn immunoreactivity, and neuromelanin in the SN**

To quantify TH cell bodies, an in-house script was developed that first identified and then counted all cells (with minimum area 100 μm2), then selected only the ones that were positive to TH. Subsequently, TH cell bodies were subtracted from the annotation, and neuromelanin-containing cells were identified with a pixel classifier with its natural brown color, made into objects, counted and subtracted from the annotation. Lastly, the script determined the %area load of TH threads with a pixel classifier. Lewy bodies were detected using cell detection based on optical density sum and specified with the following classifiers: a diameter over 6 µm, circularity, and signal intensity. Subsequently, Lewy bodies were extracted from the ROI to further quantify Lewy neurite using a pixel classifier (Supplementary Fig. 1).

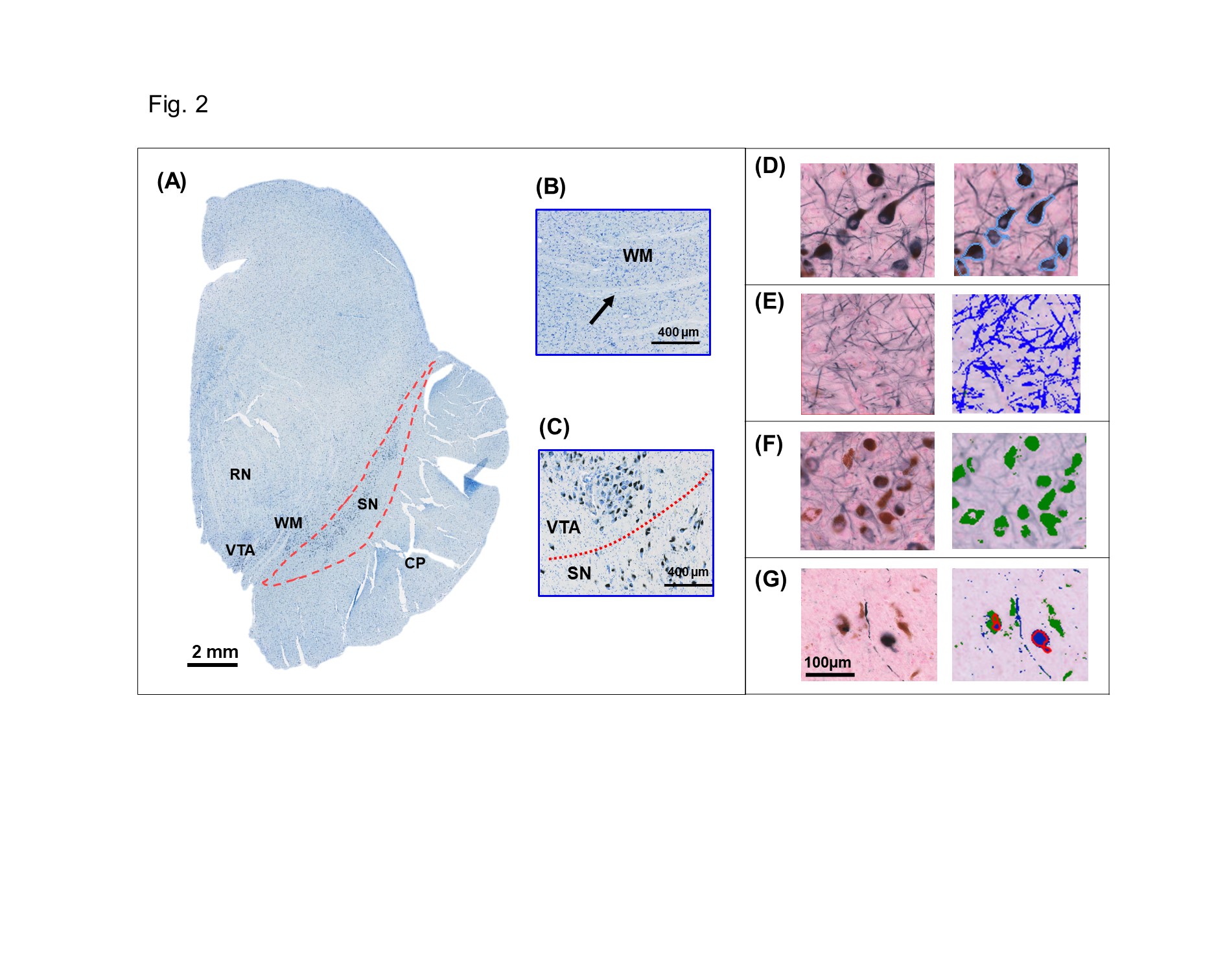
**Supplementary Fig. 1. SN delineation and quantification.** The SN was delineated on a Nissl staining, where the ventral tegmental area (VTA) served as the medial border; the red nucleus (RN) and white matter (WM) tracts served as the dorsal border and the cerebral peduncle (CP) was used as the ventral outline of the SN (A-C). Digital quantification of TH neurons (D), TH threads (E), neuromelanin (F), Lewy body and neurite (G) with in-house developed scripts. SN, substantia nigra; RN, red nucleus; VTA, ventral tegmentum area; WM, white matter; CP, cerebral peduncle.

**Results**

**Supplementary Table 1. Detail donor information**

| **Case** | **Group** | **Age at death (y)** | **Gender** | **PMD (hrs)** | **Cause of death** | **Disease duration (y)** | **Braak LB stage** | **Braak NFT stage** | **Thal phase** | **ABC score** |
| --- | --- | --- | --- | --- | --- | --- | --- | --- | --- | --- |
| 1 | PD | 83 | F | 10:35 | Euthanasia | 22 | 5 | 1 | 0 | A0B1C0 |
| 2 | PD | 69 | F | 7:05 | Aspiration pneumonia, respiratory insufficiency | 15 | 6 | 2 | 2 | A1B1C0 |
| 3 | PD | 82 | F | 9:17 | Respiratory insufficiency based on aspiration pneumonia | 17 | 6 | 2 | 2 | A1B1C0 |
| 4 | PD | 78 | M | 7:15 | Euthanasia | 17 | 6 | 2 | 1 | A1B1C0 |
| 5 | PD | 92 | M | 10:10 | Myocardial infarction | 16 | 4 | 3 | 3 | A2B2C1 |
| 6 | PD | 75 | M | 4:55 | End-stage Parkinson's disease | 20 | 6 | 2 | 3 | A2B1C0 |
| 7 | PD | 78 | M | 3:30 | End-stage Parkinson's disease | 20 | 6 | 2 | 1 | A1B1C0 |
| 8 | PD | 70 | M | 6:55 | Euthanasia | 8 | 6 | 1 | 1 | A1B1C0 |
| 9 | PD | 93 | M | 10:40 | Euthanasia | 23 | 6 | 4 | 2 | A1B2C0 |
| 10 | PDD | 83 | F | 10:40 | Deterioration, end-stage Parkinson dementia | 16 | 6 | 4 | 4 | A3B2C2 |
| 11 | PDD | 94 | F | 6:50 | Femur fracture | 10 | 6 | 4 | 4 | A3B2C2 |
| 12 | PDD | 74 | F | 8:10 | Euthanasia | 12 | 6 | 2 | 3 | A2B1C0 |
| 13 | PDD | 79 | M | 9:25 | Fall on head, subarachnoidal bleeding | 21 | 6 | 2 | 3 | A2B1C1 |
| 14 | PDD | 74 | M | 8:50 | Aspiration pneumonia | 8 | 6 | 3 | 2 | A1B2C0 |
| 15 | PDD | 81 | F | 5:30 | End-stage Parkinson's dementia | 19 | 6 | 2 | 3 | A2B1C0 |
| 16 | DLB | 66 | M | 8:00 | Euthanasia | 4 | 6 | 4 | 4 | A3B2C2 |
| 17 | DLB | 72 | M | 7:10 | Euthanasia | 5 | 6 | 2 | 3 | A2B1C2 |
| 18 | DLB | 77 | M | 7:15 | End-stage dementia with Lewy bodies | 7 | 6 | 3 | 1 | A1B2C0 |
| 19 | DLB | 91 | M | 4:30 | Euthanasia | 3 | 6 | 4 | 3 | A2B2C2 |
| 20 | DLB | 86 | F | 7:00 | Dehydration, cachexia | 6 | 6 | 4 | 5 | A3B2C2 |
| 21 | Controls | 68 | M | 8:30 | Euthanasia | NA | 0 | 1 | 2 | A1 B1 C0 |
| 22 | Controls | 63 | F | 8:10 | Euthanasia | NA | 0 | 0 | 0 | A0 B0 C0 |
| 23 | Controls | 82 | M | 10:40 | COD: gastrointestinal perforation | NA | 0 | 1 | 1 | A1 B1 C0 |
| 24 | Controls | 67 | M | 7:35 | Euthanasia | NA | 0 | 2 | 1 | A1B1C0 |
| 25 | Controls | 76 | F | 7:50 | Euthanasia | NA | 0 | 1 | 2 | A1 B1 C0 |
| 26 | Controls | 67 | M | 8:10 | COD, liver cirrhosis | NA | 0 | 1 | 1 | A1 B1 C0 |
| 27 | Controls | 77 | F | 9:00 | Euthanasia | Na | 0 | 2 | 1 | A1 B1 C0 |
| 28 | Controls | 59 | F | 8:10 | Euthanasia | NA | 0 | 0 | 0 | A0 B0 C0 |
| 29 | Controls | 71 | F | 6:50 | Lung carcinoma | NA | 0 | 1 | 2 | A1 B1 C0 |
| 30 | Controls | 74 | M | 10:20 | Euthanasia | NA | 0 | 2 | 3 | A2 B1 C0 |

Abbreviations: M, male; F, female; PMD, post-mortem delay; hr, hour; min, minutes; LB, Lewy body; NFT, neurofibrillary tangle; ABC, amyloid, Braak, CERAD (Consortium to Establish a Registry for Alzheimer’s Disease); y,years.

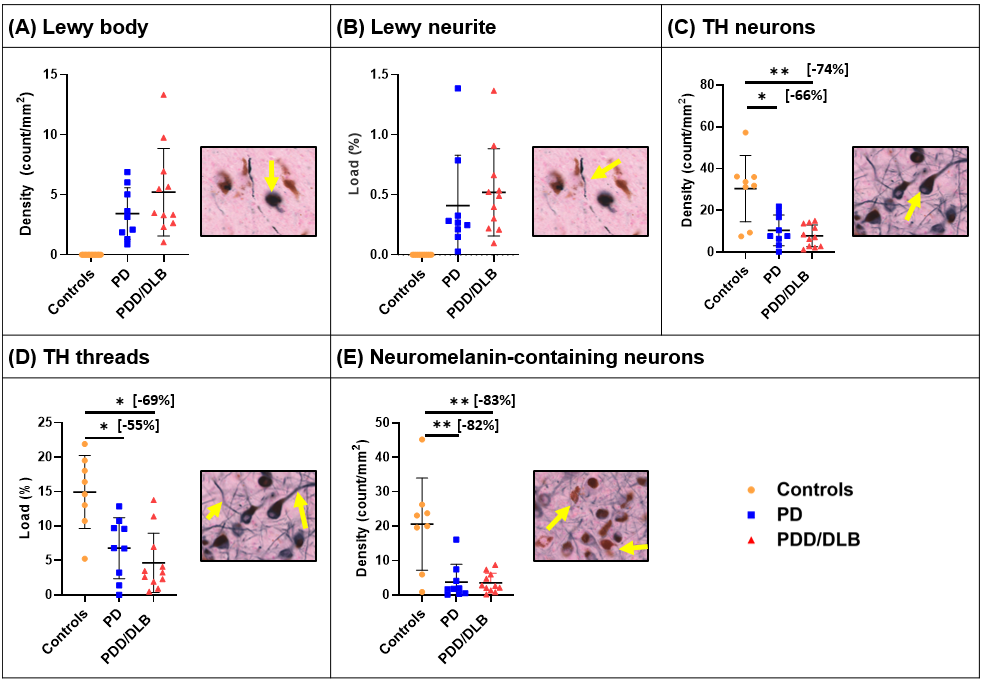

**Supplementary Fig 2. Group difference in SN histopathology.** (A) PD and PDD/DLB showed accumulation of Lewy bodies in targetoid form as indicated with the yellow arrow. (B) PD and PDD/DLB showed accumulation of Lewy neurites as line-shapes, indicated with the yellow arrow. PD and PDD/DLB did not show differences in Lewy body density nor Lewy neurite load. A comparison with controls was not made due to the biological definition of controls lacking Lewy pathology. (C) TH neurons were 66% lower in PD and 74% lower in PDD/DLB compared to controls. The yellow arrow indicates a TH cell body. (D) TH threads were 55% lower in PD and 69% lower in PDD/DLB compared to controls. The yellow arrows indicate the TH threads. (E) Neuromelanin-containing neurons were 82% lower in PD and 83 % lower in PDD/DLB compared to controls. The yellow arrows indicate the neuromelanin in its natural brown color. PD and PDD/DLB did not show difference in TH neuronal density, TH thread load and neuromelanin-containing neuronal density. Abbreviations: PD, Parkinson’s disease; PDD, Parkinson’s disease with dementia; DLB, dementia with Lewy body; TH, tyrosine hydroxylase; ns, not significant.*P<0.05 corrected, **P<0.01 corrected.

|  | **Controls** | |  | **PD+PDD/DLB** | |  | **PD** | |  | **PDD/DLB** | |  | **Compared to controls** | **PD v.s. PDD/DLB** |
| --- | --- | --- | --- | --- | --- | --- | --- | --- | --- | --- | --- | --- | --- | --- |
|  | **Mean** | **SD** |  | **Mean** | **SD** |  | **Mean** | **SD** |  | **Mean** | **SD** |  | **p-values** | **p-values** |
| **Volume**  **(mm^3^)** | 343,64 | 32,44 |  | 348,87 | 35,30 |  | 344,54 | 31,13 |  | 352,41 | 39,52 |  | Controls v.s. PD+PDD/DLB, p=.344 | *p=*.695 |
|  |  |  |  |  |  |  |  |  |  |  |  |  | Controls v.s. PD, p=.517 |  |
|  |  |  |  |  |  |  |  |  |  |  |  |  | Controls v.s. PDD/DLB, p=.314 |  |
| **FA** | 0,54 | 0,05 |  | 0,58 | 0,05 |  | 0,59 | 0,06 |  | 0,57 | 0,05 |  | Controls v.s. PD+PDD/DLB, p=.306 | *p=*.455 |
|  |  |  |  |  |  |  |  |  |  |  |  |  | Controls v.s. PD, p=.216 |  |
|  |  |  |  |  |  |  |  |  |  |  |  |  | Controls v.s. PDD/DLB, p=.524 |  |
| **MD**  **(10^-3^ mm^2^/s)** | 0,43 | 0,03 |  | 0,47 | 0,04 |  | 0,47 | 0,04 |  | 0,47 | 0,04 |  | **PD+PDD/DLB > controls, p=.048*** | *p=*.947 |
|  |  |  |  |  |  |  |  |  |  |  |  |  | Controls v.s. PD, p=.088 |  |
|  |  |  |  |  |  |  |  |  |  |  |  |  | Controls v.s. PDD/DLB, p=.070 |  |

**Supplementary Table 2. SN FA, MD and volume in each group and group differences.**

Abbreviations: PD, Parkinson’s disease; PDD, Parkinson’s disease with dementia; DLB, dementia with Lewy body; SN, substantia nigra; MD, mean diffusivity; FA, fractional anisotropy; SD, standard deviation

**Supplementary Table 3. MRI-histopathology correlations of the SN and its tracts.**

|  | **Correlations (whole cohort; PD, PDD/DLB and controls combined)** | | | | | | | |
| --- | --- | --- | --- | --- | --- | --- | --- | --- |
|  | **SN MD (10^-3^ mm^2^/s)** | |  | **Caudate tract FA** | |  | **DLPFC tract MD (10^-3^ mm^2^/s)** | |
|  | Pearson's r | p-value |  | Pearson's r | p-value |  | Pearson's r | p-value |
| **TH neurons (count/mm^2^)** | -0,273 | 0,093 |  | **-0,516** | **0,025*** |  | -0,137 | 0,524 |
| **TH threads**  **(%)** | -0,194 | 0,176 |  | -0,336 | 0,101 |  | -0,337 | 0,108 |
| **Neuromelanin-containing neurons (count/mm^2^)** | -0,094 | 0,327 |  | **-0,424** | **0,026*** |  | -0,290 | 0,169 |
|  | **Correlations (only PD and PDD/DLB combined)** | | | | | | | |
|  | **SN MD (10^-3^ mm^2^/s)** | |  | **Caudate tract FA** | |  | **DLPFC tract MD (10^-3^ mm^2^/s)** | |
|  | Pearson's r | p-value |  | Pearson's r | p-value |  | Pearson's r | p-value |
| **TH neurons (count/mm^2^)** | **-0,455** | **0,038*** |  | **-0,468** | **0,034#** |  | -0,347 | 0,094 |
| **TH threads**  **(%)** | -0,276 | 0,150 |  | **-0,438** | **0,045#** |  | -0,321 | 0,113 |
| **Neuromelanin-containing neurons (count/mm^2^)** | **-0,507** | **0,022*** |  | 0,253 | 0,172 |  | 0,186 | 0,245 |
| **Lewy bodies (count/mm^2^)** | -0,175 | 0,259 |  | 0,093 | 0,365 |  | 0,361 | 0,085 |
| **Lewy neurites (%)** | -0,142 | 0,300 |  | 0,208 | 0,220 |  | **0,501** | **0,024#** |

Abbreviations: PD, Parkinson’s disease; PDD, Parkinson’s disease with dementia; DLB, dementia with Lewy body; SN, substantia nigra; MD, mean diffusivity; FA, fractional anisotropy; DLPFC, dorsolateral prefrontal cortex; TH, tyrosine hydroxylase. *P<.05 corrected, #P<.05 uncorrected.

**References**

1.	Jonkman LE, Graaf YG de, Bulk M, et al. Normal Aging Brain Collection Amsterdam (NABCA): A comprehensive collection of postmortem high-field imaging, neuropathological and morphometric datasets of non-neurological controls. *NeuroImage Clin*. 2019;22. doi:10.1016/j.nicl.2019.101698

2.	Emre M, Aarsland D, Brown R, et al. Clinical diagnostic criteria for dementia associated with Parkinson’s disease. *Mov Disord*. 2007;22(12). doi:10.1002/mds.21507
